# A randomized placebo-controlled trial on the antidepressant effects of the psychedelic ayahuasca in treatment-resistant depression

**DOI:** 10.1101/103531

**Authors:** Fernanda Palhano-Fontes, Dayanna Barreto, Heloisa Onias, Katia C Andrade, Morgana Novaes, Jessica A Pessoa, Sergio A Mota-Rolim, Flavia Osório, Rafael Sanches, Rafael G dos Santos, Luís F Tófoli, Gabriela de Oliveira Silveira, Mauricio Yonamine, Jordi Riba, Francisco RR Santos, Antonio A Silva-Junior, João Alchieri, Nicole L Galvão-Coelho, Bruno Lobão-Soares, Jaime Hallak, Emerson Arcoverde, João P Maia-de-Oliveira, Draulio B Araújo

## Abstract

Recent open label trials show that psychedelics, such as ayahuasca, hold promise as fast-onset antidepressants in treatment-resistant depression. In order to further test the antidepressant effects of ayahuasca, we conducted a parallel-arm, double-blind randomized placebo-controlled trial in 29 patients with treatment-resistant depression. Patients received a single dose of either ayahuasca or placebo. Changes in depression severity were assessed with the Montgomery–Åsberg Depression Rating Scale (MADRS) and the Hamilton Depression Rating scale (HAM-D). Assessments were made at baseline, and at one (D1), two (D2) and seven (D7) days after dosing. We observed significant antidepressant effects of ayahuasca when compared to placebo at all timepoints. MADRS scores were significantly lower in the ayahuasca group compared to placebo (at D1 and D2: p=0.04; and at D7: p<0.0001). Between-group effect sizes increased from D1 to D7 (D1: Cohen’ s d=0.84; D2: Cohen’ s d=0.84; D7: Cohen’ s d=1.49). Response rates were high for both groups at D1 and D2, and significantly higher in the ayahuasca group at D7 (64% vs. 27%; p=0.04), while remission rate was marginally significant at D7 (36% vs. 7%, p=0.054). To our knowledge, this is the first controlled trial to test a psychedelic substance in treatment-resistant depression. Overall, this study brings new evidence supporting the safety and therapeutic value of ayahuasca, dosed within an appropriate setting, to help treat depression.

## Introduction

The World Health Organization estimates that 350 million people suffer from depression, and about one-third do not respond to appropriate courses of at least three antidepressants.^1–3^ Most currently available antidepressants have a similar efficacy profile and mechanisms of action, based on the modulation of brain monoamines, and take about two weeks to start being effective.^1–3^

Recent evidence, however, shows a rapid and significant antidepressant effect of ketamine, an Nmethyl-D-aspartate (NMDA) antagonist frequently used in anesthesia.^4–7^ Namely, in randomized placebocontrolled trials, the antidepressant effects of ketamine in treatment-resistant depression peaked one day after dosing and remained significant for about seven days.^4–7^

Additionally, research with serotonergic psychedelics has gained momentum.^8^ A few centers around the world are currently exploring how these substances affect the brain, and also probing their potential in treating different psychiatric conditions, including mood disorders.^9–13^ For instance, recent open label trials show that psychedelics, such as ayahuasca and psilocybin, hold promise as fast-onset antidepressants in treatment-resistant patients.^10,13^ Moreover, randomized controlled trials have recently shown that psilocybin reduces anxiety and depression symptoms in patients with lifethreatening cancer.^11,12^

Ayahuasca is a brew traditionally used for healing and spiritual purposes by indigenous populations of the Amazon Basin. In the 1930s, it began to be used in religious settings of Brazilian small urban centers, reaching large cities in the 1980s and expanding since then to several other parts of the world.^14^ In Brazil, ayahuasca has a legal status for ritual use since 1987. Ayahuasca is prepared by decoction of two plants: *Psychotria viridis* that contains the psychedelic N,N-dimethyltryptamine (N,N-DMT), a serotonin and sigma-1 receptors agonist,^15^ and *Banisteriopsis caapi*, rich in reversible monoamine oxidase inhibitors such as harmine, harmaline, and tetrahydroharmine.^16^

The acute psychological effects of ayahuasca last around 4h and include intense perceptual, cognitive and affective changes.^16–18^ Although nausea, vomiting and diarrhea are often reported, mounting evidence points to a positive safety profile of ayahuasca. For instance, ayahuasca is not addictive and has not been associated with psychopathological, personality or cognitive deterioration, and it promotes only moderate sympathomimetic effects.^19–21^ The main concern is rare instances of prolonged increases in psychotomimetic symptoms, especially in individuals prone to psychosis.^18,22^

In a recent open label trial, 17 patients with major depressive disorder attended a single dosing session with ayahuasca. Depression severity was assessed before, during and after dosing by the Hamilton Depression Rating Scale (HAM-D) and the Montgomery– Åsberg Depression Rating Scale (MADRS).^10^ Significant reduction in depression severity was found already in the first hours after dosing, an effect that remained significant for 21 days.^10,23^

Although promising, these studies have not controlled for the placebo effect, which can be remarkably high in clinical trials for depression, reaching 30-40% of the patients.^24^ To address this issue, and to further test the antidepressant effects of ayahuasca, we conducted a placebo-controlled trial in patients with treatment-resistant major depression.

## Materials and Methods

### Study design and participants

This study is a double-blind parallel-arm randomized placebo-controlled trial. Patients were recruited from psychiatrist referrals from local outpatient psychiatric units or through media advertisements. All procedures took place at the Onofre Lopes University Hospital (HUOL), Natal-RN, Brazil. The University Hospital Research Ethics Committee approved the study (# 579.479), and all subjects provided written informed consent before participation.

We recruited adults aged 18-60 years who met criteria for unipolar major depressive disorder as diagnosed by the Structured Clinical Interview for Axis I (DSM-IV). Only treatment-resistant patients were selected, defined herein as those with inadequate responses to at least two antidepressant medications from different classes.^2^ Selected patients were in a current depressive episode of moderate-to-severe at screening (HAM-D≥17). We adopted the following exclusion criteria: previous experience with ayahuasca, current medical disease based on history, pregnancy, current or previous history of neurological disorders, history of schizophrenia or bipolar affective disorder, history of mania or hypomania, use of substances of abuse, and suicidal risk.

### Randomization and masking

Patients were randomly assigned (1:1) to receive ayahuasca or placebo, using permuted blocks of size 10. All investigators and patients were blind to intervention assignment, which was kept only in the database and with the pharmacy administrators. Masking was further achieved by ensuring that all patients were naïve to ayahuasca, and by randomly assigning a different psychiatrist for patient follow-up assessments. Both ayahuasca and placebo were kept in identical amber glass bottles.

### Procedures

The substance used as placebo was a liquid designed to simulate the organoleptic properties of ayahuasca, such as a bitter and sour taste, and a brownish color. It contained water, yeast, citric acid, zinc sulphate and a caramel colorant. The presence of zinc sulphate also produced low to modest gastrointestinal distress. A single ayahuasca batch was used throughout the study. The batch was prepared and provided free of charge by a branch of the Barquinha church based in the city of Ji-Paraná-RO, Brazil.

To assess alkaloids concentrations and stability of the batch, samples of ayahuasca were quantified at two different moments by mass spectroscopy analysis (details in suppl. material). On average, the ayahuasca contained (mean±SD): 0.36±0.01 mg/mL of N,N-DMT, 1.86±0.11 mg/mL of harmine, 0.24±0.03 mg/mL of harmaline, and 1.20±0.05 mg/mL of tetrahydroharmine. Data from the individual assessments are shown in supplementary table S1.

After screening, patients underwent a washout period adjusted to the half-life time of their current antidepressant medication, which typically lasted two weeks. During the dosing session, patients were not under any antidepressant medication, and a new treatment scheme was introduced only seven days after dosing. If needed, benzodiazepines were allowed as a supporting hypnotic and/or anxiolytic agents (suppl. table S2).

Two clinical scales for depression assessed changes in depression severity: MADRS and HAM-D. Assessments were made at baseline (one day before dosing), and at one (D1), two (D2) and seven (D7) days after dosing. HAM-D was applied only at baseline and D7, as it was designed to access depression symptoms present in the last week.^25^

Dosing sessions lasted approximately 8 hours, from 8:00 to 16:00, and intake usually occurred at 10:00. After a light breakfast, patients were reminded about the effects they could experience, and strategies to help alleviating eventual difficulties. Sessions took place in a quiet and comfortable living room-like environment, with a bed, a recliner, controlled temperature, natural and dimmed light.

Patients received a single dose of 1 ml/kg of placebo or ayahuasca adjusted to contain 0.36 mg/kg of N,N-DMT. They were asked to remain quiet, with their eyes closed, while focusing on their body, thoughts and emotions. They were also allowed to listen to a predefined music playlist. Patients received support throughout the session from at least two investigators who remained in a room next door, offering assistance when needed. Acute effects were assessed with the Clinician-Administered Dissociative States Scale (CADSS) and the Brief Psychiatric Rating Scale (BPRS), which were applied at -10 min, +1:40h, +2:40h and +4:00h after intake.

When acute psychedelic effects ceased, patients debriefed their experience, and had a final psychiatric evaluation. Around 16:00 they could go home accompanied by a relative or friend. Patients were asked to return for follow-up assessments one, two and seven days after dosing, when also a new antidepressant was prescribed.

### Outcomes

The primary outcome measure was the change in depression severity assessed by the HAM-D scale, comparing baseline with D7. In this study we chose D7 as the primary assessment point to allow direct comparison with previous randomized trials of ketamine in treatment-resistant depression.^4–7^ Furthermore, D7 was chosen to avoid interaction with the new antidepressant medication, which was prescribed also at D7. The secondary outcome was the change in MADRS scores from baseline to D1, D2 and D7. We examined the proportion of patients meeting response criteria, defined as a reduction of 50% or more in baseline scores. Remission rates, defined as HAM-D≤7 or MADRS≤10, were also examined. Both response and remission rates were computed using scores from HAM-D (at D7) and MADRS (at D1, D2, and D7). Safety and tolerability was also assessed by evaluation of dissociative and psychotomimetic symptoms using the CADSS and BPRS–Positive Subscale (BPRS+) during the dosing session.

### Statistical analysis

Analyses adhered to a modified intent-to-treat principle, including all patients who completed assessments at baseline, dosing and D7. An estimated sample size of 42 patients is needed to provide 80% power to detect a 5-point HAM-D difference (standardized effect size=0.9) for differences of ayahuasca and placebo between baseline and D7 with two-sided 5% significance. A fixed-effects linear mixed model, with baseline scores as covariate, examined changes in HAM-D scores at D7, and in MADRS scores at D1, D2 and D7. A Toeplitz covariance structure was the best fit to the data according to Akaike’ s information criterion. Missing data were estimated using restricted maximum likelihood estimation. Main effects and treatment vs. time interaction were evaluated. Post-hoc t-tests were performed for between-groups comparisons at all timepoints, and Sidak’ s test was used to control for multiple comparisons. Significance was evaluated at p<0.05, two-tailed. Cohen’ s d effect sizes were obtained for between and with-in group comparisons. Between-group effect sizes were calculated using the estimated means of each group at each timepoint. For within-group comparisons, effect sizes of each treatment were calculated separately, using the differences between an endpoint and baseline values. Differences in proportion of responders/non-responders and remitters/non-remitters between treatments were estimated using Fisher’ s exact test. Odds ratio (OR) and number needed to treat (NNT) were also calculated. Data from patients whose HAM-D or MADRS scores reduced by 50% or more between washout onset and baseline assessment, or were in remission in the day of dosing, were not considered for statistical analysis. We used IBM SPSS Statistic 20 and Prism 7 to run the analyses. This study is registered with http://clinicaltrials.gov (NCT02914769).

## Results

From January 2014 to June 2016, we assessed 218 patients for eligibility, and 35 met criteria for the trial. Six volunteers had to be excluded: five no longer met criteria for depression in the day of dosing, and one dropped out before dosing. Data from 29 patients were included in the analysis: 14 in the ayahuasca group and 15 in the placebo group. Figure 1 shows the trial profile.

**Figure 1.**
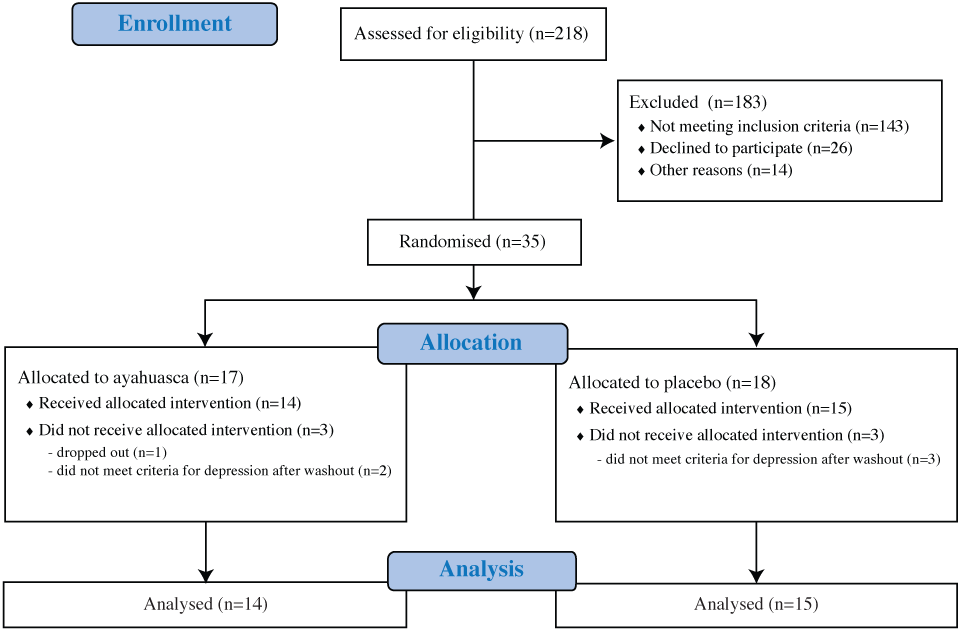
Trial profile.

On average, patients met criteria for moderateto-severe depression (mean±SD): HAM-D=21.83±5.35; MADRS=33.03±6.49. They had been experiencing depressive symptoms for 11.03±9.70 years, and had tried 3.86±1.66 different previous unsuccessful antidepressants. Two patients had previous history of electroconvulsive therapy (ECT). Most patients (76%) had a comorbid personality disorder, and 31% had comorbid anxiety disorder. All patients were under regular use of benzodiazepines during the trial (suppl. table S2).

Demographic and clinical characteristics are summarized in table 1 (suppl. table S2 for individual data). All patients were Brazilian, mostly female (72%) adults (42.03±11.66 yo) from low socioeconomic status backgrounds: low educated (41% with <8 years of formal education) and living in low household income (41% earn <2 minimum wages).

**Table 1.** Socio-demographic & clinical characteristics

|  | Ayahuasca | Placebo |
| --- | --- | --- |
| Participants, n | 14 | 15 |
| Age (years) | 39.71±11.26 | 44.2±11.98 |
| Gender (M/F) | 3/11 | 5/10 |
| Unemployed (%) | 7/14 (50) | 8/15 (53) |
| Household income |  |  |
| < 2 minimum wages (%) | 6/14 (43) | 6/15 (40) |
| 2-5 wages (%) | 4/14 (28) | 7/15 (47) |
| 6-10 wages (%) | 1/14 (7) | 1/15 (6.6) |
| 11 or more wages (%) | 3/14 (21) | 1/15 (6.6) |
| Education |  |  |
| Up to 8 years, n (%) | 6/14 (43) | 6/15 (40) |
| 9-11 years, n (%) | 3/14 (21) | 5/15 (33) |
| 12-16 years, n (%) | 2/14 (14) | 2/15 (13) |
| 17 or more years, n (%) | 3/14 (21) | 2/15 (13) |
| Religion (%) |  |  |
| Catholic | 7/14 (50) | 5/15 (33) |
| Protestant | 4/14 (28) | 1/15 (6.6) |
| Other | 0/14 (0) | 4/15 (27) |
| No religion | 3/14 (21) | 5/15 (33) |
| Ethnicity (%) |  |  |
| Caucasian | 9/14 (64) | 8/15 (54) |
| Black | 1/14 (7) | 0/15 (0) |
| Pardo | 4/14 (28) | 7/15 (47) |
| **Clinical characteristics** |  |  |
| Age of depression onset (years) | 30.93±10.19 | 30.87±13.39 |
| Illness duration (years) | 8.78±6.25 | 13.13±11.92 |
| Number of previous episodes | 2.71±1.32 | 3.53±1.76 |
| Length of current episode (months) | 14.71±18.92 | 10.13±9.15 |
| Failed antidepressant medications | 3.93±1.44 | 3.8±1.89 |
| History of ECT (%) | 1/14 (7) | 1/15 (6.6) |
| History of psychotherapy (%) | 11/14 (79) | 12/15 (80) |
| Anxiety disorder (%) | 5/14 (36) | 5/15 (33) |
| Personality disorder (%) | 10/14 (71) | 12/15 (80) |
| Melancholic (%) | 12/14 (83) | 12/15 (80) |
| Atypical (%) | 2/14 (14) | 3/15 (20) |
| Baseline HAM-D | 24.07±5.34 | 19.73±4.59 |
| Baseline MADRS | 36.14±6.12 | 30.13±5.55 |

Values are (mean ± SD); M=male; F=female; ECT=electroconvulsive therapy.

Figure 2 shows changes in HAM-D scores from baseline to seven days after dosing. We observe a significant between-groups difference at D7 (F_1_=6.31; p=0.019) with patients treated with ayahuasca presenting less severity when compared to patients treated with placebo (suppl. figure S1 for individual scores). Between-group effect size is large at D7 (Cohen’ s d=0.98; CI 95%: 0.21 to 1.75). Within-group effect size (suppl. table S3) is large for the ayahuasca group (Cohen’ s d=2.22; CI 95%: 1.28 to 3.17), and medium for the placebo (Cohen’ s d=0.46; CI 95%: -0.27 to 1.18).

**Figure 2.**
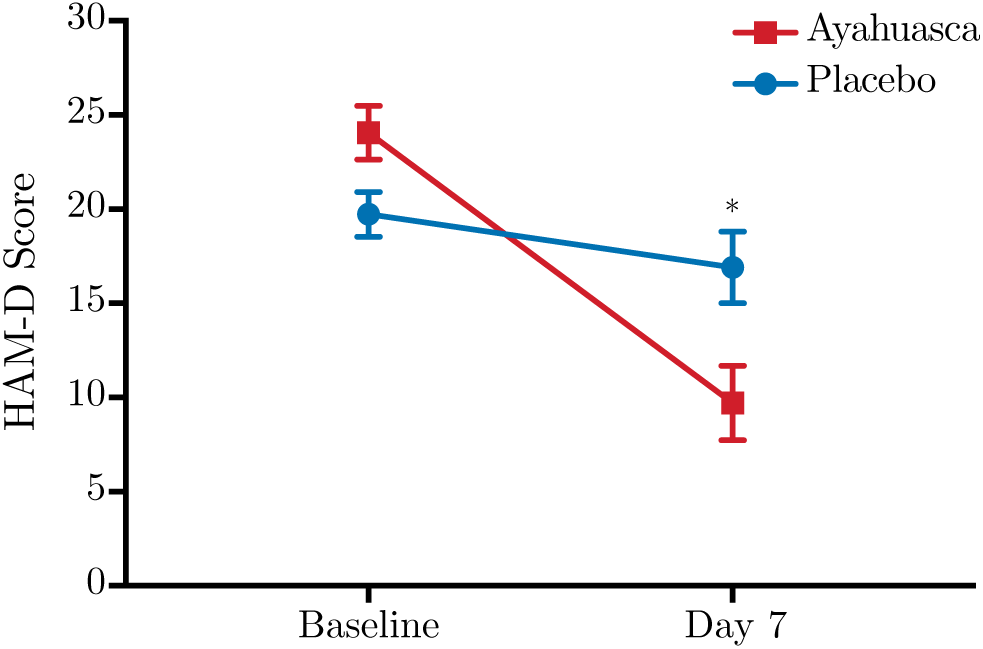
HAM-D scores at baseline and seven days after dosing. Statistical analysis shows a significant difference between ayahuasca (red squares) and placebo (blue circles) seven days after dosing (p=0.019). Between-group effect size is high (Cohen’ s d=0.98). Values are (mean ± SEM). HAM-D scores: mild depression (8–16), moderate (17–23), severe (≥ 24).

Figure 3 shows mean MADRS scores as a function of time. Linear mixed model shows a significant effect for time (F_2,34.4_=3.96; p=0.028), treatment (F_1,27.7_=10.52; p=0.003), but no treatment vs. time interaction (F_2,34.4_=1.77; p=0.185). Individual scores are presented in supplementary figure S2. Post-hoc analysis shows a significant difference between groups at D1 (F_1,49.7_=4.58; p=0.04), D2 (F_1,50.3_=4.67; p=0.04) and D7 (F_1,47_=14.81; p<0.0001).

**Figure 3.**
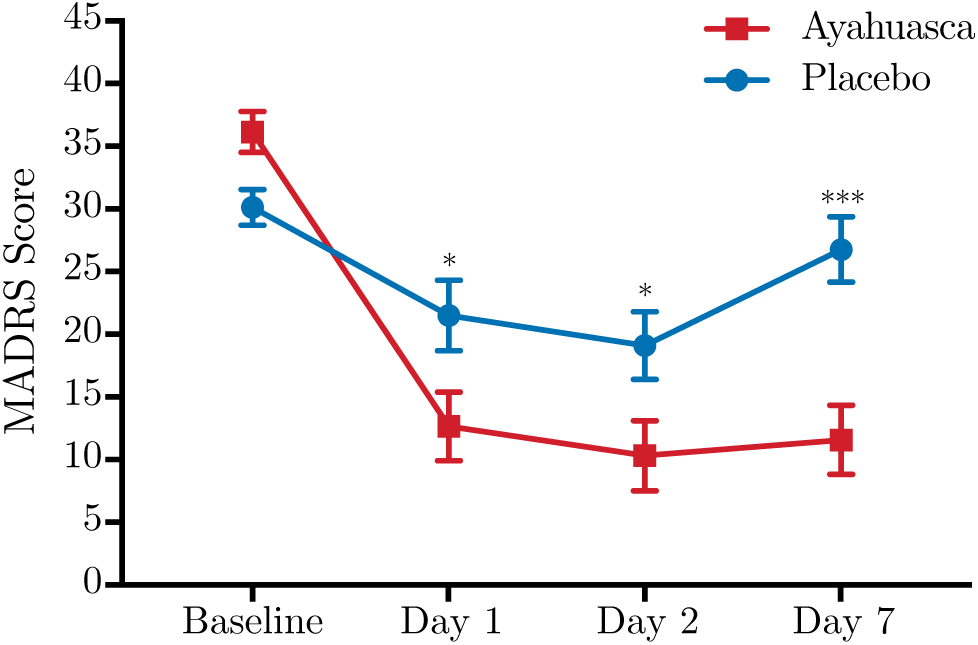
MADRS scores as a function of time. Significant differences are observed between ayahuasca (red squares) and placebo (blue circles) at D1 (p=0.04), D2 (p=0.04) and D7 (p<0.0001). Between groups effect sizes are high at all timepoints after dosing: D1 (Cohen’ s d=0.84), D2 (Cohen’ s d=0.84) and D7 (Cohen’ s d=1.49). Values are (mean ± SEM). MADRS scores: mild depression (11–19), moderate (20–34), severe (≥ 35). *p <0.05; ***p <0.0001.

Between-groups effect size is large at D1 (Cohen’ s d=0.84; CI 95%: 0.05 to 1.62) and D2 (Cohen’ s d=0.84; CI 95%: 0.05 to 1.63) and largest at D7 (Cohen’ s d=1.49; CI 95%: 0.67 to 2.32). Within-group effect sizes (suppl. table S4) are large for the ayahuasca group at all timepoints after dosing (Cohen’ s d=2.78, CI 95%: 1.74 to 3.82 at D1, 3.05, CI 95%: 1.94 to 4.16 at D2, 2.90, CI 95%: 1.84 to 3.97 at D7).

HAM-D response rate at D7 was significantly different between-groups, with 57% of responders in the ayahuasca group against 20% in the placebo group (OR=5.33 [95% CI: 1.11 to 22.58]; p=0.04; NNT=2.69). The difference HAM-D remission rate was marginally significant at D7: 43% in ayahuasca vs. 13% in placebo (OR=4.87 [95% CI: 0.77 to 26.73]; p=0.07; NNT=3.39).

Figure 4a shows the MADRS response rates as a function of time. At D1, response rates were high for both groups: 50% in the ayahuasca group, and 46% in the placebo group (OR=1.17 [95% CI: 0.26 to 5.48]; p=0.87; NNT=26). At D2, they remained high in both groups: 77% in the ayahuasca group and 64% in the placebo (OR=1.85 [95% CI: 0.29 to 8.40]; p=0.43; NNT=7.91). Response rate was statistically different at D7: 64% of responders in the ayahuasca group, and 27% in the placebo (OR=4.95 [95% CI: 1.11 to 21.02]; p=0.04; NNT=2.66).

**Figure 2.**
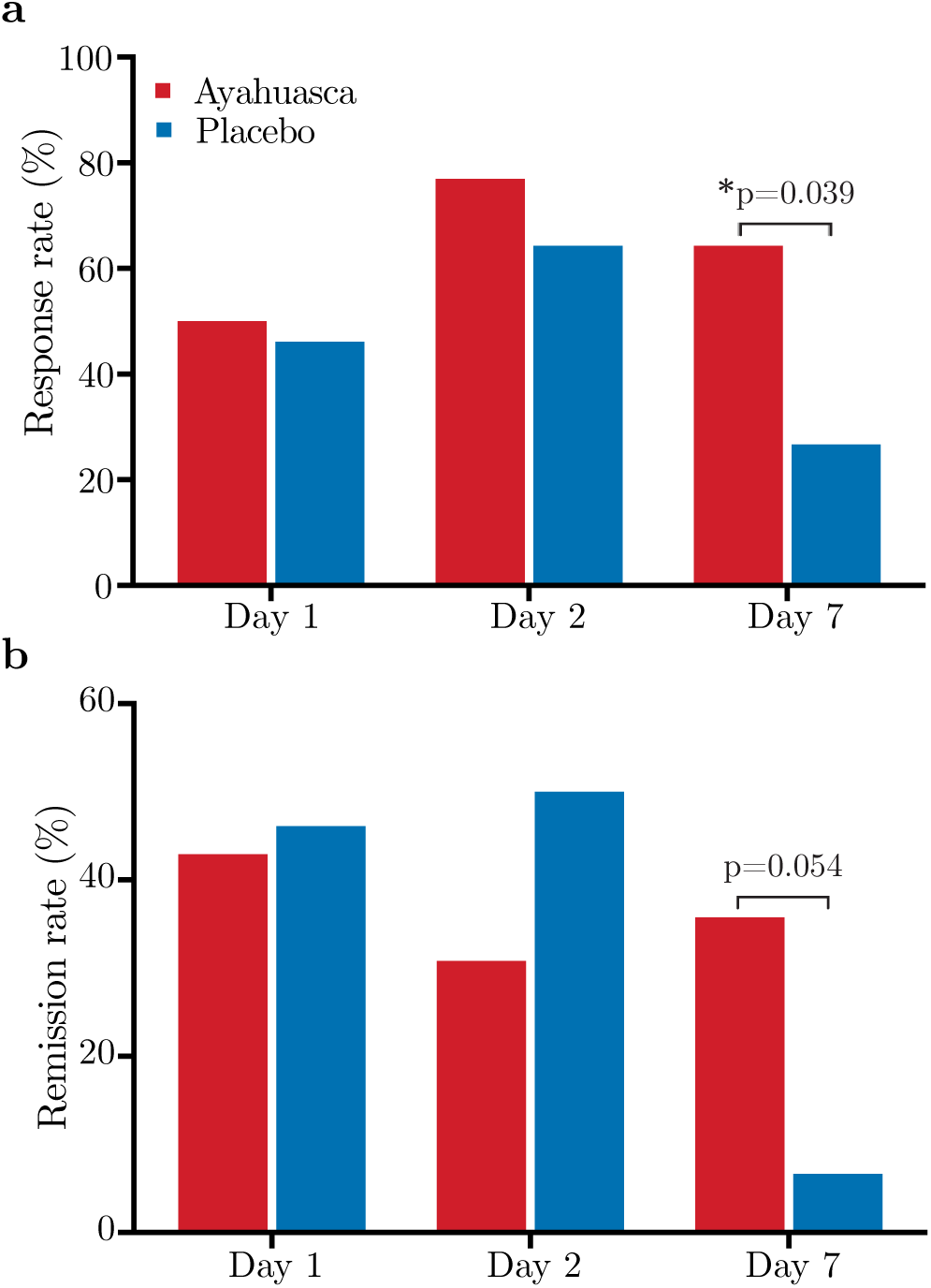
Response and remission rates as a function of time. Response **(a)** and remission **(b)** rates were high for both groups at D1 and D2. At D7, responses rate is significantly higher for ayahuasca (OR=4.95 [95% CI: 1.11 to 21.02]; p=0.04; NNT=2.66), while remission rate is marginally significant (OR=7.78 [95% CI: 0.81 to 77.48]; p=0.054; NNT=3.44).

Figure 4b shows the MADRS remission rates as a function of time. At D1 and D2, remission rates were not statistically different between groups (p=0.86 and p=0.31, respectively). At D7 MADRS remission rate was marginally significant: 36% of remitters in the ayahuasca group and 7% in the placebo (OR=7.78 [95% CI: 0.81 to 77.48]; p=0.054; NNT=3.44). Supplementary figure S2 shows individual MADRS scores %-changes from baseline, at all timepoints. Although individual variance is high, overall, we find decreased scores for all subjects in the ayahuasca group.

The most frequently observed adverse effects in the ayahuasca group included nausea (71%), vomiting (57%), transient anxiety (50%), transient headache (42%), and restlessness (50%) (suppl. table S5). Patients exhibited transient dissociative and psychotomimetic symptoms as measured by CADSS and BPRS+ scales, with slightly increased scores +1h40 after ayahuasca intake: 34.8% (BPRS+) and 21.6% (CADSS) (suppl. table S6).

## Discussion

We found evidence for rapid antidepressant effects after a single dosing session with ayahuasca, when compared to placebo. Depression severity changed significantly but differently for the ayahuasca and placebo groups. At all timepoints before dosing, improvements in the psychiatric scales observed in the ayahuasca group were significantly higher than those of the placebo group, with increasing between-group effect sizes from D1 to D7. Response rates were high for both groups at D1 and D2, and were significantly higher in the ayahuasca group at D7. Remission rate was marginally significant at D7.

The within-group effect size found for ayahuasca at D7 (Cohen’ s d=2.22) is compatible with our earlier open label study (Cohen’ s d at D7=1.83),^10^ and compatible with the one found in a recent open label trial with psilocybin for depression (Hedges’ g=3.1).^13^

Our results are comparable with randomized controlled trials that used ketamine in treatment-resistant depression. ^4–7^ Although both ketamine and ayahuasca are associated with rapid antidepressant effects, their response time-courses and mechanisms of action seem to differ. Previous studies with ketamine have found the largest between-group effect size at D1 (Cohen’ s d=0.89), reducing towards D7 (Cohen’ s d=0.41).^4–7^ In contrast, the effect sizes observed herein were smallest at D1 (Cohen’ s d=0.84), and largest at D7 (Cohen’ s d=1.49). These differences are also reflected in the response rate. At D1, the response rate to ketamine lies between 37-70%, whereas in our study 50% of the patients responded to ayahuasca. At D7, the ketamine response rate ranges between 7-35%,^4–7^ while in our study 64% responded to ayahuasca.

The placebo effect was high in our study, and higher than most studies with ketamine.^4–7^ While we find a response rate to placebo of 46% at D1, and 26% at D7, ketamine trials have found a placebo effect on the order of 0-6% at D1, and 0-11% at D7.^4–7^ Several factors may contribute to the high placebo effects observed here. First, higher placebo effects are found in patients with low socioeconomic status,^24^ which was the case of our study, where most of the patients were living under significant psychosocial stressors. During our trial, on the other hand, they stayed at a very comfortable and very supportive environment. It is more of a caring effect, than properly a placebo effect. Second, patients with comorbid personality disorders present higher placebo responses.^25^ In our study, most patients (76%) also suffered from personality disorders, most of them in cluster B.

A growing body of evidence gives support to the observed rapid antidepressant effects of ayahuasca. For instance, the sigma-1 receptor (σ1R) has recently been implicated in depression, and was reported to be activated by N,N-DMT.^1,15^ Moreover, it has been shown that the administration of σ1R agonists results in antidepressant-like effects, which are blocked by σ1R antagonism.^1^ Furthermore, σ1R upregulates neurotrophic factors such as brain derived neurotrophic factor (BDNF) and nerve growth factor (NGF), proteins whose regulation and expression seem to be involved in the pathophysiology of depression.^1^

Studies in animal models reported that chronic administration of harmine reduces immobility time, increases climbing and swimming time, reverses anhedonia, increases adrenal gland weight, and increases BDNF levels in the hippocampus.^26,27^ All of these are compatible with antidepressant effects. Likewise, harmine seems to stimulate neurogenesis of human neural progenitor cells, derived from pluripotent stem cells, a mechanism also observed in rodents following antidepressant treatment.^28^ Also, a recent study in rodents found that a single ayahuasca dose increases swimming time in a forced-swim test.^29^

Over the last two decades, mental health evaluations of regular ayahuasca consumers have shown preserved cognitive function, increased well-being, reduction of anxiety, and depressive symptoms when compared to non-ayahuasca consumers.^19,20^ Moreover, a recent study observed that a single dose of ayahuasca enhanced mindfulness-related capacities,^30^ and meditation practices have been associated with antidepressant effects.^31^

Brain circuits modulated by psychedelics show great overlap with those involved in mood disorders.^8^ We recently found that a single ayahuasca session in patients with depression increases blood flow in brain regions consistently implicated in the regulation of mood and emotions,^3^ such as the left nucleus accumbens, right insula and left subgenual area.^10^ Moreover, we have shown that ayahuasca reduces the activity of the Default Mode Network (DMN),^32^ a brain network found to be hyperactive in depression, possibly due to rumination.^3,33^

No serious adverse effects were observed during or after the dosing session. Although 100% of the patients reported feeling safe during the ayahuasca session, it was not necessarily a pleasant experience. In fact, some patients reported the opposite, as the experience was accompanied by much psychological distress. Most patients reported nausea, and about 57% have vomited, although vomiting is traditionally not considered a side effect of ayahuasca, but rather part of a purging process.^17^

Although promising, this study has some caveats and limitations worth mentioning. The number of participants is modest, and therefore randomized trials in larger populations are necessary. The study was limited to patients with treatment-resistant depression, with long course of illness, and high comorbid personality disorder, which altogether precludes a simple extension of these results to other classes of depression. Another challenge of the research with psychedelics is maintaining double blindness, as the effects of psychedelics are unique. We were particularly keen to ensure blindness throughout the entire experiment, and to that end we adopted a series of additional measures to preserve blindness. All patients were naïve to ayahuasca, with no previous experience with any other psychedelic substance. Clinical evaluations involved a team of five psychiatrists. For every patient, one psychiatrist was responsible for clinical evaluation during the dosing session, and another for the follow-up assessments. The substance used as placebo increased anxiety and induced nausea. In fact, five patients misclassified placebo as ayahuasca, and two of them showed response at D7. Therefore, we believe blindness was adequately preserved in our study.

Since the prohibition of psychedelics in the late 1960s, research with these substances has almost come to a halt. Before research restrictions, psychedelics were at early stage testing for many psychiatric conditions, including obsessive-compulsive disorder and alcohol dependence. By mid 1960s, over 40.000 subjects had participated in clinical research with psychedelics, most of them in uncontrolled settings.^8^ To our knowledge, this is the first randomized placebo-controlled trial to investigate the antidepressant potential of a psychedelic in a population of patients with treatment-resistant depression. Overall, this study brings new evidence supporting the safety and therapeutic value of psychedelics, dosed within an appropriate setting, to help treat depression.

## Acknowledgments

The authors would like to express their gratitude to all patients who volunteered for this experiment. To the Brain Institute and to the Hospital Universitário Onofre Lopes (HUOL), both from the Federal University of Rio Grande do Norte (UFRN) for always giving the necessary institutional support. To the laboratory of Pharmacognosy for helping with placebo preparation. To Edilsom Fernandes, for carefully preparing the Ayahuasca batch used in our study, and for the fruitful discussion, particularly about the dosing session. To Sidarta Ribeiro, for the enthusiastic support throughout the study. To Dr. Ricardo Lagreca, for the unconditional support. To Beatriz Labate, for the fruitful discussions. To Altay Souza and João Sato for assistance with statistical analysis. To Deborah Maia, Ranna Brito, Tayrine Lopes, Lízie Brasileiro, Artur Morais, Isaac Campos, Brígida Albuquerque, Marianna Lucena, Fernanda Araújo, Raíssa Nóbrega, Marina Leonardo, Kaique Andrade, Rodolfo Lira, Giuliana Travassos, for helping with data acquisition. To Prof. Octávio Pontes-Neto and Adriano Tort for critical review of the manuscript. To the Brazilian federal funding agencies CNPq (grants #466760/2014 & #479466/2013) and CAPES (grants #1677/2012 & #1577/2013) for providing financial support.

